# A novel PSII photosynthetic control is activated in anoxic cultures of green algae

**DOI:** 10.1101/2021.12.01.470728

**Authors:** Yuval Milrad, Valéria Nagy, Szilvia Z. Tóth, Iftach Yacoby

## Abstract

Photosynthetic green algae face an ever-changing environment of fluctuating light as well as unstable oxygen levels, which via the production of free radicals constantly challenges the integrity of the photosynthetic complexes. To face such challenges, a complex photosynthetic control network monitors and tightly control the membrane redox potential. Here, we show that not only that the photosynthetic control set the rate limiting step of photosynthetic linear electron flow, but also, upon its ultimate dissipation, it triggers intrinsic alternations in the activity of the photosynthetic complexes. These changes have a grave and prolonged effect on the activity of photosystem II, leading to a massive 3-fold decrease in its electron output. We came into this conclusion via studying a variety of green algae species and applying advance mass-spectrometry and diverse spectroscopic techniques. Our results shed new light on the mechanism of photosynthetic regulation, and provide new target for improving photosynthesis.

## Introduction

Photosynthetic electron flow is crucial for the development of complex life on earth, as in this process sunlight is captured as a primary energy source for most living organisms. This process is mostly renowned for its role in O_2_ evolution by photosystem II (PSII) of cyanobacteria, algae and plants (Antal et al., 2020; Vinyard et al., 2013). In this process, two water molecules are split to generate one O_2_ molecule by the oxygen evolving complex (OEC) (Peloquin and Britt, 2001; Umena et al., 2011). Electrons are then channeled via a stable plastoquinone site (Q_A_) to mobile plastoquinone or semi-quinone, which is located on the acceptor site of PSII (Q_B_). The protonation of the plastoquinone in Q_B_ site is mediated via a non-heme iron, which is regulated by the binding of a HCO_3_^-^ molecule (Saito et al., 2013; Tikhonov et al., 2018). Once plastoquinone is fully reduced to generate plastoquinol, it is released from the Q_B_ site and diffuses into the thylakoid membrane, where it predominantly reduces cytochrome *b*_*6*_*f* (Cyt*b*_*6*_*f*), which channels electrons to downstream metabolites via photosystem I (PSI). The process of electron flow is constantly regulated to bypass barriers according to the availability of metabolites of the photosynthetic apparatus. Redox poise and sub-localization of complexes in the thylakoid membrane were both postulated to pose “photosynthetic control” and maintain electron flux to fit the system’s capacity (Kirchhoff et al., 2017; Schöttler and Tóth, 2014). Green microalgae, which are continuously subjected to environmental changes, have evolved several regulatory mechanisms to cope with fast transitions, such as: chlororespiration by plastid terminal oxidase (PTOX) (Houille-vernes et al., 2011; Nawrocki et al., 2015), which oxidizes plastoquinol, NADPH oxidation by flavodiiron proteins (FLVs) (Burlacot et al., 2018; Chaux et al., 2017; Jokel et al., 2018) and the formation of superoxide radicals via the Mehler reaction in PSI (Kozuleva et al., 2021). These processes enable cells to cope with fast transitions from darkness as they increase the amount of available downstream products of both photosystems, and thus alleviate acceptor side limitations. However, when a transition is taking place under strict anaerobiosis, due to the cells’ respiration or external O_2_ scavenging, H_2_ evolution by hydrogenase, which is otherwise prone to inactivation by O_2_ (Stripp et al., 2009), remains the only effective valve to cope with an excess of energy upon such sudden light exposure (Godaux et al., 2015; Milrad et al., 2018).

The kinetics of H_2_ production stands in a direct competition with NADP^+^ reduction by ferredoxin-NADP^+^ reductase (FNR) (Kosourov et al., 2020; Milrad et al., 2018), a crucial reactant for carbohydrates synthesis via the Calvin-Benson-Bassham cycle (CBB). At light onset, H_2_ is evolved at a high rate, while CO_2_ fixation (by the CBB cycle) is hindered (Allen, 2003). Then, as the imbalanced ratio of NADPH and ATP is alleviated, and CO_2_ fixation is activated, H_2_ evolution rates gradually decline (Ghysels et al., 2013; Godaux et al., 2015). To bypass such limitations, it was previously shown that challenging the cells with fluctuating light, ranging minutes or below, can limit O_2_ to low levels and thus improve the sustainability of H_2_ production (Jokel et al., 2019; Kosourov et al., 2018; Milrad et al., 2018). However, such attempts resolved another, yet unidentified barrier for H_2_ evolution; while the initial exposure to light following an hour of dark anoxia triggers a fast flux of electrons, as reported by high rates of H_2_ evolution, in successive exposures, under minutes range of dark anaerobiosis, H_2_ evolution rate is drastically reduced (by 3-folds), regardless of the number of fluctuations (Milrad et al., 2018). To date, the mechanism responsible for this dramatic decline remains unresolved. In this work, we explored the origins of this massive decrease via the assessments of global and local electron fluxes in intact algal cultures. We recorded and integrated the electron transport processes from OEC to processes downstream of PSI. Our results suggest that the redox activated “photosynthetic control” regulates the activity of PSII and cause a massive reduction in its electron output.

## Results

### Following a short light exposure, photosynthetic productivity is decreased by 3-fold

Dark anoxic cultures of green microalgae immediately emit H_2_ at high rates upon exposure to light. To examine the kinetics of this H_2_ evolution, we studied several species of green algae: *Chlamydomonas reinhardtii* (Cr), *Chlorella sorokiniana* (Cs) and a newly isolated strain of *Monorophidium* (Mo), from the Yarkon River in Tel-Aviv, Israel. Each species was cultivated in mixotrophic medium and incubated under dark anoxia for an hour. Following incubation, we exposed the cells to light (370 µE m^−2^ sec^−1^) for two minutes, and the concentrations of diffused H_2_ and O_2_ were monitored using a membrane inlet mass spectrometer (MIMS) (Liran et al., 2016) (**Figure. 1**). In all tested strains, following an initial burst, H_2_ evolution rate declined rapidly to a complete cessation. As H_2_ accumulation plateaued; the light was turned off. Importantly, tracking the O_2_ accumulation enabled us to keep the cells under darkness for enough time to completely respire O_2_ (3-5 minutes), hence maintain anoxia. We then illuminated the cells again, for two minutes and observed that H_2_ evolution resumed. These light/dark fluctuations were repeated four times, and although H_2_ re-evolved in all successive light exposures, we observed a ∼3-fold global decrease in H_2_ accumulation rates in comparison with the initial exposure to light in all tested species.

**Figure 1:**
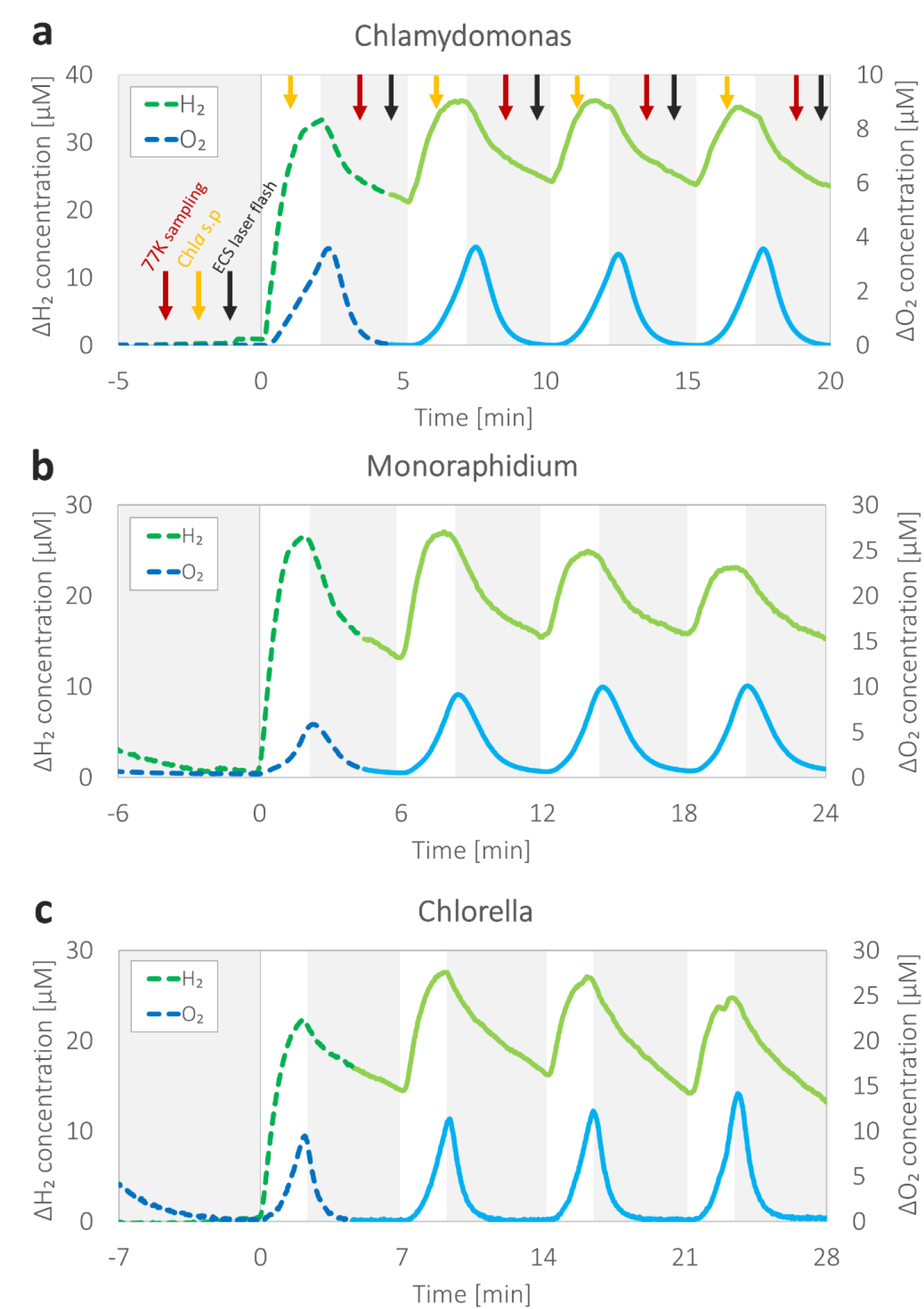
Hydrogen and oxygen production following light and dark fluctuations. Following an hour of dark anaerobic incubation, three algal species: *Chlamydomonas reinhardtii* (**a**, Cr), a newly isolated strain of *Monoraphidium* (**b**, Mo) and *Chlorella sorokiniana* (**c**, Cs), were examined in a MIMS for H_2_ (green) and O_2_ (blue) accumulation. The species were challenged with light fluctuations of 2 minutes under illumination, at irradiances of 370, 250 and 1000 µE m^-2^ s^-1^ for Cr, Mo, and Cs respectively, followed by either 3, 4 or 5 minutes of darkness for Cr, Mo, and Cs, respectively. In all species, the initial exposure to light (dashed) triggered a 3-fold rise in H_2_ accumulation compared to successive exposures (solid). Several additional measurements (marked by arrows) were conducted for Cr: (Yellow)-determination of maximal Chl*a* fluorescence by a saturating pulse, (red)-cell sampling for 77K fluorescence and (black)-determination of the ECS absorbance changes that where induced by charge separations following a single 5 ns laser flash. The ECS signal was measured using a Joliot type spectrophotometer (JTS-100) equipped with a BiLED (520-546 nm) measuring lamp. The results of these additional measurements will be presented and discussed further on. Each curve represents the averaged result of at least three biological repetitions.

To further study this phenomenon, we focused our research on a wild type strain of Cr, CC124 (137c mt-). Since all successive light exposures showed the same kinetics of H_2_ evolution, they were united to an average successive trace and compared *versus* the initial light exposure (**Figure. 2**). We observed that in each light exposure, H_2_ production initiated at high rates (**Figure. 2a**), which declined as electrons were diverted toward other competing processes (Milrad et al., 2018). To test whether these differences are limited only to mixotrophic growth, we conducted a similar test, using cells which were cultivated under autotrophic conditions (**Supp. 1**), and observed the same decline in H_2_ evolution rates (between initial and successive fluctuations). The kinetics of O_2_ net accumulation were also altered, as its rate increased upon successive exposures. Interestingly, we also observed that the initial exposure triggered an immediate linear increase in net O_2_ evolution; however, this trait was not observed in autotrophic conditions (**Supp. 1**) or other algal species (**Figure. 2b**) and could therefore be attributed to differences in acetate assimilation. CO_2_ concentrations were also monitored, and observed to rapidly accumulate at light onset in both initial and successive light exposures. Such evolution has been the subject of much research, and is attributed to either a reverse activity of carbonic anhydrase, which is part of the carbon concentrating mechanism (CCM) (Kaplan and Reinhold, 1999; Milrad et al., 2021), and/or the activity of PSII itself, as was recently proposed (Brinkert et al., 2016; Koroidov et al., 2014; Kosourov et al., 2020; Shevela et al., 2020). To compare the kinetics of CO_2_ evolution/fixation between initial and successive light fluctuations, we calculated the rates of CO_2_ concentration change (**Figure. 2c**). We observed that the increased rate of CO_2_ emission at light onset was the same between initial and successive exposures. However, the time needed for the flux to start declining toward net CO_2_ fixation was longer in successive light exposures (16 seconds in the initial light exposure compared to 35 seconds in successive fluctuations, seen as a positive rate peak in **Figure. 2c**). This could be a result of either a delayed activation of the CBB cycle in successive exposures or a prolonged release of CO_2_. To gain better perspective on the activation of the CBB cycle, we simultaneously monitored the concentration of NADPH by measuring its fluorescence (**Figure. 2d**). It was previously shown that under anoxia, at light onset, NADPH levels decline due to its oxidation via Type-II NADPH dehydrogenase (Nda2) and/or shuttling via the Malate dehydrogenase (MDH) (Milrad et al., 2021). Our results here show that the rapid decline of NADPH concentration reaches steady state much faster in the initial exposure, approximately 10 seconds in the initial light exposure *versus* a minute in the successive fluctuations. This indicates that a balanced redox poise, which is a prerequisite for activation of the CBB cycle, is achieved faster in the initial exposure.

**Figure 2:**
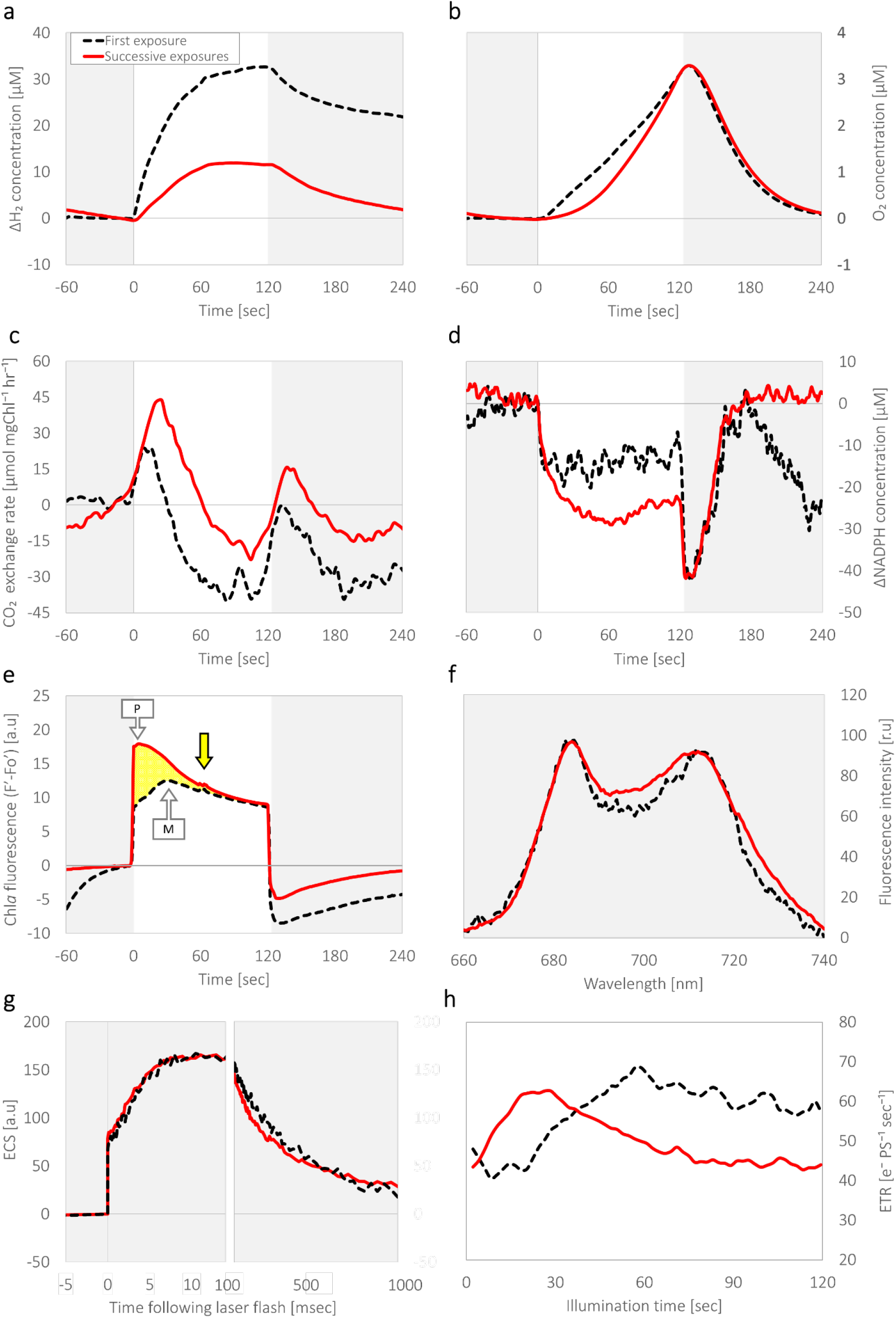
Electron transfer shifts under fluctuating light. *C. reinhardtii* wild-type strain CC124 cells were incubated for an hour under dark anaerobiosis, after which they were challenged with light fluctuations of 2 minutes under illumination (at an irradiance of 370 µE m^-2^ s^-1^, white background), followed by 3 minutes of darkness (gray background, as shown in Figure 1). Shown are the differences between the initial light exposure (dashed black) and the average of the successive exposures (solid red). H_2_ (**a**) and O_2_ (**b**) concentrations were measured using MIMS, in addition to CO_2_ for which the exchange rates were calculated (**c**). Simultaneously, the changes in NADPH concentration were also measured using Dual-PAM-100 equipped with DUAL-ENADPH (**d**), in addition to Chl*a* fluorescence (**e**). During illuminations, the cells were exposed to saturating pulses (marked with an arrow in **e**, see also **Figure. 1a**), and the traces were normalized to Fm’ to compare the kinetics of the changes in PSII photosynthetic efficiency. Yellow area in panel (**e**) mark the observed differences between Chl*a* fluorescence traces, which were induced by light fluctuations. Prior to and in-between illuminations, we sampled the cells (as displayed in **Figure. 1a**, see red arrows) and immediately tested their fluorescence spectra under 77K (**f**). Charge separation was evaluated via electrochromic shift (ECS) measurements (**g**), for which the cells were exposed to a laser flash 30 seconds prior to each light exposure (as displayed in **Figure. 1a**, see black arrows). To assess the overall electron transfer rate (**h**), the cells were exposed to short intervals of darkness (60 msec), during illumination (every 2 seconds) and the rate of ECS decline was determined. Each curve represents the averaged result of at least three biological repetitions.

### Linear electron flow is down regulated by light exposure

Net O_2_ measurements are not sufficient to correctly estimate the quantum yield of PSII because they can only reflect its gross production at some level. Therefore, we also tested alterations in photosynthetic efficiency by measuring chlorophyll *a* (Chl*a*) fluorescence (Schreiber et al., 1995; Sipka et al., 2021) (**Figure. 2e)**. Due to the “Kautsky effect”, the slow kinetics of Chl*a* fluorescence (termed here as F’), following dark anoxia features two peaks (Stirbet and Govindjee, 2011). The first peak “P”, is reached when all Q_B_ sites are occupied with plastoquinol (Strasserf et al., 1995). The second peak “M” is reached due to either; i) the activation of downstream processes, specifically CO_2_ fixation by the CBB cycle (Dufková et al., 2019), ii) conformational changes in PSII (Krause, 1988) or iii) distribution of LHCII antenna complexes (state transition) (Kodru et al., 2015). Following “P” and “M” peaks, F’ reaches steady state.

As expected, we observed that at the initial light exposure, F’ increased for a duration of 30 seconds (dashed in **Figure. 2e**) (Tolleter et al., 2011). In contrast, Successive illuminations triggered an immediate sharp increase in F’ (in the timeframe of P) and featured no additional peak. It should be noted however that following 60 seconds of illumination all fluorescence traces equalized, and steadily declined in the same manner. To assess whether heat dissipation or state transition were altered, we exposed the cells to a saturating pulse, as the traces reached steady state (see yellow arrows in **Figures. 1a and 2e**), and determined the maximal fluorescence (Fm’). We observed no significant changes between light exposures (**Supp. 2**), indicating minimal changes if any in the magnitude of non-photochemical quenching, nor a decrease of PSII efficiency due to state transition as the cells reach steady state. To verify that indeed no shifts in the localization of LHCII antennas took place, we sampled cells directly from the experiment’s cuvette (see red arrows in **Figure. 1a)** and examined their fluorescence spectra at 77°K (**Figure. 2f**). We observed that the peaks, which are observed at ∼680nm and ∼710nm, show no alternations between the first and consecutive light pulses, which indicates that the time of light exposure (2 minutes) is not sufficient to generate state transition (Takahashi et al., 2013), which is in accordance with previous research (Ghysels et al., 2013).

To verify that changes are not originating from alternations in the overall efficiency of the photosystems, we examined differences in electrochromic shifts (ECS) between light exposures, by tracking changes in their absorbance (Bailleul et al., 2010) (of 520-546 nm) (**Figure. 2g**, see also; black arrows in **Figure. 1a**). When cells are exposed to a single turnover laser flash their absorbance is changed via a 3-phase shift (Bailleul et al., 2010). The initial increase in the ECS signal (termed “phase a”) lasts less than a millisecond and is a product of charge separations taking place in both photosystems, hence decreased signal can report on their degradation. During the second phase (termed “phase b”), which usually lasts up to 10 milliseconds (Buchert et al., 2020), ECS signal increases due to the build-up of membrane potential, mainly via Cyt*b*_*6*_*f*. In the final phase (termed “phase c”), the signal decreases exponentially due to the dissipation of the membrane potential via ATPase activity. Here we observed that all flashes, which were given before each light exposure, triggered an identical shift, indicating that the redox state at light onset was similar between light fluctuations (**Figure, 2g**). We then measured the general electron flow during light fluctuations by examining short dark interval relaxation kinetics (DIRK) during illumination (Sacksteder and Kramer, 2000)_’_(Bailleul et al., 2010). The results (**Figure, 2h**) show that upon illumination, following a prolonged dark anaerobic incubation, the general electron flux is stable on a relatively low rate (∼40 e^−^ PS ^−1^ sec ^−1^) for 20 seconds before it starts increasing. The flux rate then stabilizes following a minute of illumination at a rate of ∼60 e^−^ PS ^−1^ sec ^−1^. Interestingly, the rate of electron flux during successive light exposures starts also at ∼40 e^−^ PS ^−1^ sec ^−1^. However, it does not delay and as soon as the cells are exposed to light a rapid increase is observed. Following 30 seconds of illumination, the electron flux rate fades until it resumes its original value of ∼40 e^−^ PS ^−1^ sec ^−1^.

### Light activation of photosynthetic regulation mechanisms

Notably, the measurements at successive illuminations are puzzling because while the global electron flow is larger, the apparent quantum yield of PSII and Hydrogenase activity are both lower. To solve this discrepancy, we monitored the redox state of P700^+^ (PSI) (**Figure. 3)** according to (Klughammer and Schreiber, 1994). The results show a gradual increase in the difference of PSI oxidation between light and dark steady states (Ps’-Po’) along all light exposures, which is saturated following a minute of illumination at approximately the same value. Interestingly, in successive light exposures, the value of Ps’-Po’ is quite high at light onset, suggesting that the cells are already donor side limited, *i*.*e*., less electrons arrive at PSI. Next, we determined the differences in between Pm’-Ps’ values (related to the apparent quantum yield of PSI (Buchert et al., 2020)). Interestingly, we observed that at the initial light exposure, the apparent quantum yield of PSI is slowly increased, in contrast to the successive exposure, where it is already optimal at light onset. Taken together, the results of global electron flow and P700^+^ are in line with previous report (Godaux et al., 2015), which showed that at first light onset, PSI cyclic electron flow is gradually increased for 2 minutes. This enhanced activity increases the proton motive force across the thylakoid membrane, which was postulated to initiate “photosynthetic control” (Schöttler et al., 2015), and thus cause a shift in the rate limiting steps of the photosynthetic apparatus. Photosynthetic control is activated by elevated redox poise, which is dependent on the duration of irradiance. Such regulation was proposed to diminish linear electron flow (Schöttler and Tóth, 2014), and subsequently decrease H_2_ production. Hence, one could suggest that such changes in electron flow should require sufficient time for a build-up of the redox poise across the thylakoid membrane. To test this hypothesis, following dark incubation of an hour, we exposed the cultures to light, for a varied duration of time, between 5-180 seconds (**Figure. 4**). Then, the cells were kept under darkness for three minutes, before they were exposed to light for a second time, for additional two minutes. We then tracked the evolution of H_2_ (**Figure. 4a**) and slow kinetics of Chl*a* fluorescence (**Figure. 4b**) during the second 2 minutes illumination, and plotted the results in accordance with the preceding duration of the initial exposure (**Figure.4)**. Our results show that the activation of the photosynthetic regulation mechanism is taking place after minimal time of ∼20 seconds. In addition, we monitored the build-up of redox poise, which is associated with the activation of “photosynthetic control”, by measuring changes in the kinetics of ECS build-up following an exposure to laser flash (**Figure. 4c**). We observed that, although no differences in rapid charge separation at “phase a” are apparent (**Supp. 3**), the buildup of the redox potential, seen by the lack of an apparent rise in the trace at “phase b”, is gradual which is in accordance with the decrease in hydrogenase activity. Such changes of the redox potential should also decrease the electron flux via Cyt*b*_*6*_*f*, due to the activation of “photosynthetic control”, and therefore gradually pose a donor-side limitation on PSI and an acceptor side limitation on PSII. Such limitation could ultimately result in a decreased rate of linear electron flow, which is also seen as a lower rate of H_2_ production.

**Figure 3:**
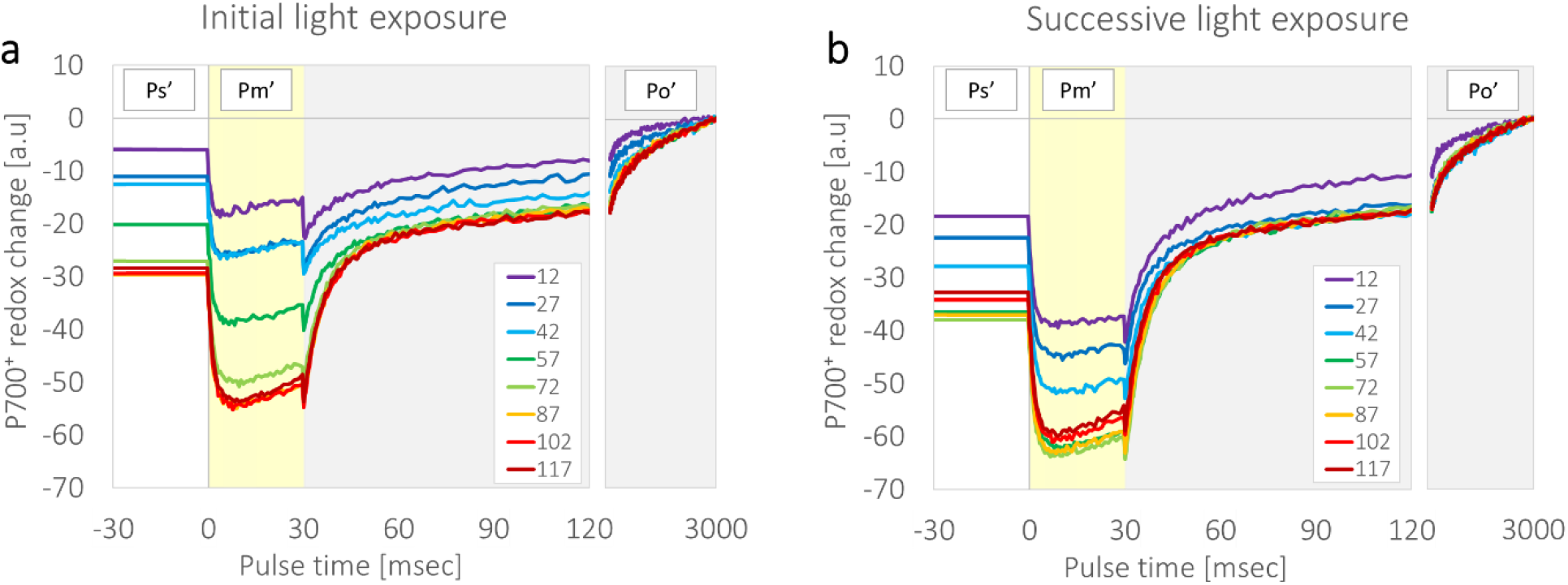
P700+ reduction under fluctuating light exposures. *C. reinhardtii* wild-type strain CC124 cells were incubated for an hour under dark anaerobiosis, after which they were exposed to variable timing of actinic light (220 µE m ^−2^ sec^−1^ white background; The Color index of the plots starts with purple - following 12 seconds of actinic light to dark red - following 117 seconds of actinic light). P700^+^ (PSI) redox state was measured during continuous actinic light (Ps’), after which the cells were exposed to a high light pulse (30 msec at 2000 µE m ^−2^ sec ^−1^, Pm’, yellow background), followed by 3 seconds of darkness (Po’=0, at dark steady state, gray background). These high light pulses were shot every 15 seconds. Regardless when [Ps’] was measured, the total duration of the initial light exposure (**a**) was set to 2 minutes, after which the light was turned off for 3 minutes before resuming the measurement again for a successive light exposure (**b**). Graphs represent the averaged results of three biological repeats.

**Figure 4:**
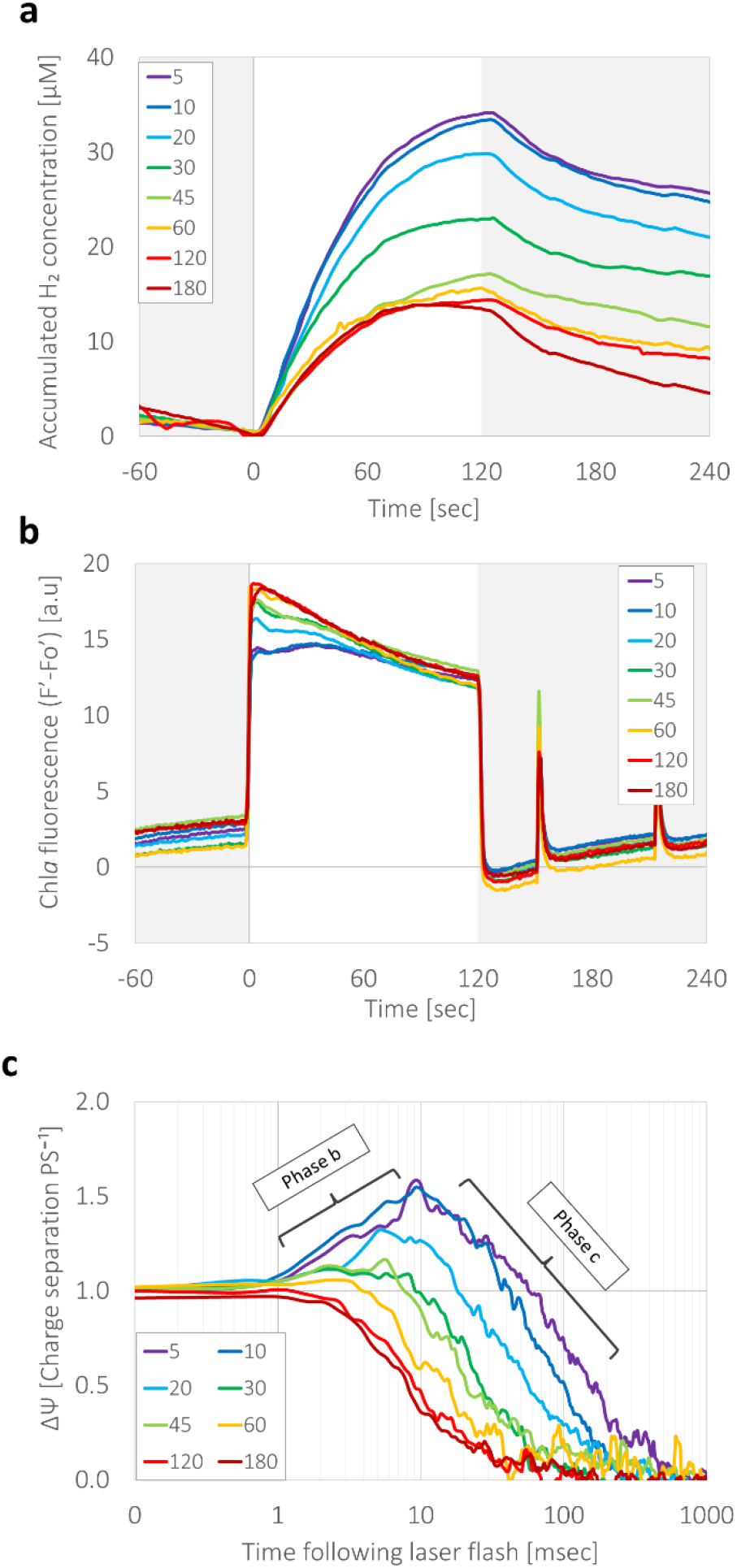
Light duration of photosynthetic control activation. **(a)** *C. reinhardtii* wild-type strain CC124 cells were incubated for an hour under dark anaerobiosis, after which they were illuminated (370 µE m^−2^ sec ^−1^) for a duration of either 5, 10, 20, 30, 45, 60, 120 or 180 seconds. Following that, the cells were kept in darkness for 3 minutes before they were illuminated again for 2 minutes, during which, H_2_ accumulation (**a**) and Chl*a* fluorescence (**b**) were measured. Three seconds following the initial light exposure, the cells were exposed to a 5 ns laser flash, and charge separation was measured (**c**). Color index in all panels matches the duration of the initial light exposure (ranging seconds): 5-purple, 10-blue, 20-cyan, 30-green, 45-light green, 60-yellow, 120-red and 180-dark red. Each experiment was repeated using at least three biological replicates.

### Dark recovery of the initial electron transfer rate

Since we observed that light exposure triggered a decrease in linear electron flow, we examined whether such modulations are permanent or could be reversed due to prolonged darkness. To that aim, we gradually increased the duration of darkness in between light exposures, which were set to a constant two minutes of light exposure (**Supp. 4**). Dark exposures were timed to: 3, 5, 7, 10 and 15 minutes, having an additional 3 minutes of darkness before the last illumination to verify that the observed alternations are indeed due to the duration of darkness, rather than the total time that passed in the experiment (marked with an asterisk; 3*). In addition, to eliminate possible interferences by accumulated O_2_, we added an O_2_ scavenger, glucose oxidase (GOx, supplied with glucose and catalase (Roberty et al., 2014)). The results show that the decrease in H_2_ evolution, following 3 minutes of darkness is not affected by the absence of O_2_ (**Figure. 5a**), and that H_2_ evolution rate increases in accordance with prolonged darkness. We observed that the cells require at least 15 minutes of darkness to fully regain the initial 3-fold faster H_2_ evolution rate. Since we postulated that this decrease is correlated to the activity of PSII, we added the inhibitors; 3-(3,4-dichlorophenyl)-1,1-dimethylurea (DCMU) and hydroxylamine (HA) to the cells, prior to illumination (**Figure. 5b**). The results show that the rate of H_2_ accumulation is impaired in all light fluctuations, following a short burst of H_2_ evolution at light onset, as was previously described (Milrad et al., 2021). In addition, we observed no significant changes in-between fluctuations. To assess whether the build-up of membranal potential was also affected by the blocked PSII, we evaluated the differences in charge separation in the absence (**Figure. 5c**) or presence (**Figure. 5d**) of DCMU and HA. To our surprise, “phase a” of the ECS signal showed no decrease due to the presence of DCMU and HA (**Supp. 5**), allowing us to normalize the results as electrons per PS as shown earlier. Interestingly, the addition of DCMU and HA shortened the time needed for retrieving the original rise in “phase b” to less than two minutes under darkness.

**Figure. Dark time: Figure 5:**
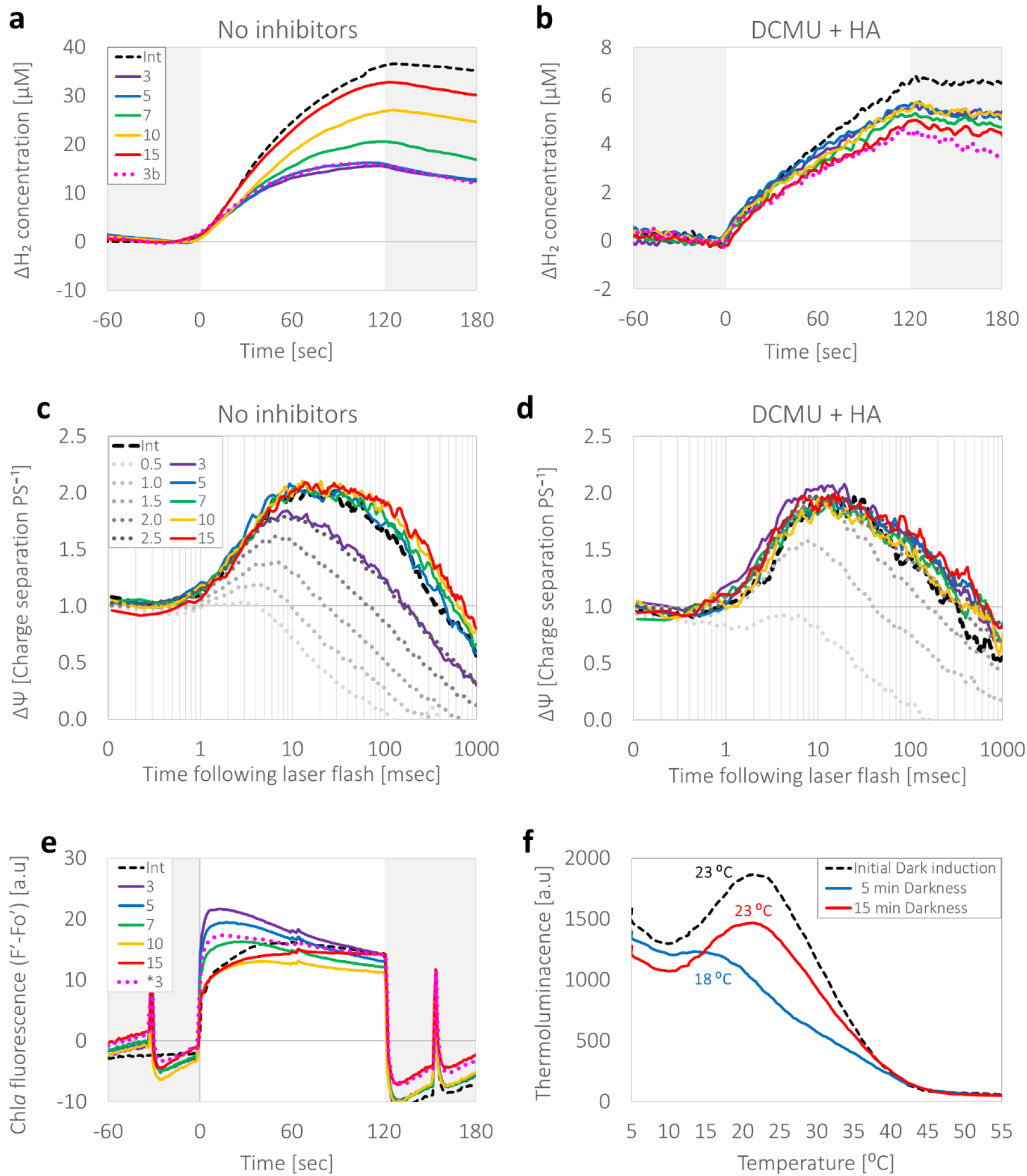
Darkness reestablish the initial fast electron flux. Following an hour of dark incubation in the presence of O_2_ scavengers (GOx), *C. reinhardtii* wild-type strain CC124 cells were subjected to a series of light exposures (370 µE m^-2^ s^-1^ for 2 minutes, white background) hatched with an increasing duration of dark incubations 0-15 minutes. H_2_ accumulation was detected in the absence (**a**) or presence (**b**) of DCMU and hydroxylamine (HA). Presented are the measured concentration of H_2_ according to each preceded dark incubation. Charge separation of a laser flash induced cells were measured prior (Int) and 0.5-15 minutes following two minutes of light exposure, in either the absence (**c**) or presence (**d**) of DCMU and HA. (**e**) Chl*a* fluorescence of the untreated cells was measured, simultaneously to H_2_ evolution as presented in panel a. During each illumination, the cells were exposed to a saturating pulse and maximal fluorescence (Fm’) was calculated. Basal fluorescence (Fo’) was subtracted from the measured fluorescence (Ft), and the results were plotted relative to Fm’. (**f**) Cells’ thermoluminescence was measured after 2 single turnover flashes (STF). The cells were tested following an hour of dark anaerobic incubation (Int). Samples were then illuminated for 2 minutes and dark-adapted for either 5 (blue) or 15 (red) minutes. In all panels, the color index is marked by the duration of preceding darkness, where the initial exposure (Int) is marked in dashed black, short flashes following: 0.5, 1, 1.5, 2, and 2.5 minutes of darkness are marked in gradient dotted gray, and following: 3, 5, 7, 10, and 15 minutes of darkness are marked in: purple, blue, green, yellow and red respectively. In panels a, b and e, the measurements following an additional 3 minutes of darkness at the end of the experiment are marked in dotted pink. Each experiment was repeated using at least three biological replicates. Error bars indicate standard error (n ≥ 3).

Since we previously observed alternations in PSII apparent activity, we tracked changes in Chl*a* fluorescence (**Figure. 5e**), simultaneously with the MIMS measurements presented earlier for the untreated cells (**Figure. 5a**). We observed the same gradual alternations in the early rise of the trace for ∼60 seconds (**Figure. 5e**). To verify that state transition did not lower the fluorescence signal, we exposed the cells to a saturating pulse a minute prior and 30 seconds after each illumination and observed no differences between paired maximal fluorescence values (**Supp. 6**). We then examined alternations in the internal electron transport chain of PSII, by measuring the effect of light exposure on cells’ Thermoluminescence (TL). Light emission from a pre-illuminated sample during temperature increase provides valuable information about the charge recombination reactions occurring within PSII (for a review, see (Ducruet, 2013; Ducruet and Vass, 2009)). Since we aimed to test the functionality of the OEC, we exposed the cells to two flashes, which result in a mixed recombination of Q_B_^-^ with S_2_ and S_3_, and therefore increase the gained signal for the “S_2/3_ B band” (Ducruet, 2013). We tested cell samples following an hour of dark anoxic incubation, and observed a peak of the “S_2/3_ B band” at a temperature of 23.2±0.2 °C (**Figure. 5f**). We then exposed the cells to two minutes of light, followed by 5 minutes of darkness and took a second sample. Interestingly, the illumination triggered a major decrease in the intensity of the B band, indicating that less charge recombination occurred between the S_2_ and S_3_ states of the OEC and Q_B_^-^ or, alternatively, recombination occurred via non-radiative pathways (Vass and Cser, 2009). In addition, the maximal value of the band was downshifted to 18.2±2.2 °C. These changes indicate that the redox equilibrium between Q_A_^-^ and Q_B_^-^ may have changed towards Q_A_^-^, facilitating charge recombination with the S-states of the donor side. Finally, to assess whether indeed these shifts are reversed during longer periods of anoxic darkness, we exposed the cells to a second two-minute period of illumination followed by 15 minutes of anoxic darkness, and indeed observed that both the intensity shift of the B band and the temperature in which it peaked resumed to their initial state following dark anoxia.

## Discussion

In nature, green algae are subjected to various environmental changes. Since they mostly rely on the operation of the photosynthetic apparatus, many of these changes are resolved by the cells’ ability to rapidly alternate the course of their metabolic electron flow (Alric et al., 2010; Hemschemeier and Happe, 2011). Indeed the activation or deactivation of electron sinks has been the topic of much research recently, and was shown to hold a crucial role in the ability of the cells to sustain functionality (Burlacot et al., 2019; Godaux et al., 2015; Jokel et al., 2019; Milrad et al., 2018). However, in this work, we show that in green algae under anoxia upstream regulations have a profound effect on the overall productivity of the apparatus, decreasing the electron flux, due to light exposure. (**Figure. 1**). Our results suggest that the decrease in the overall flux of electrons is originated from a decrease in the apparent linear electron flow from PSII, and is governed by the “photosynthetic control”.

Following anaerobic induction, hydrogenase serves as a major electron sink to energy flow downstream to PSI. Although it alleviates acceptor side limitations on PSI (Ghysels et al., 2013), it is still not as efficient as the CBB cycle, and therefore the apparent quantum yield of PSI initiates at low rates (**Figure. 3a**). Then, as cyclic electron flow generates an additional redox potential, PSI efficiency increases (**Figure. 3a**), CO_2_ fixation initiates (**Figure. 2c**), and a general increase in the electron flux appear (**Figure. 2h**). It was previously shown that the generated proton motive force activates “photosynthetic control” in which the rate of Cyt*b*_*6*_*f* activity is decreased (Schöttler and Tóth, 2014). Such decrease shifts the limitations that are posed on PSI, from its acceptor to donor side (**Figure. 3**). This would also result in continuous reduced pool of plastoquinol and hinder the activity of PSII, thus generating an excessive pressure on its acceptor side (**Figure. 2e**). Indeed, it was postulated that such pressure, could dissociate the HCO_3_^-^ molecule which is located next to the non-heme iron in PSII and trigger conformational changes in PSII. In turn, the exchange rate of the plastoquinone/ol in the Q_B_ site decreases (Brinkert et al., 2016; Koroidov et al., 2014; Shevela et al., 2020). Thus, the plastoquinone pool may function as a switch impeding O_2_ evolution and charge separation in PSII. Similar postulations were made by Cardona and coworkers, stating that: “An acceptor side switch, should it exist, could for example be triggered by the formation of Q_A_^-^ before the Q_B_H_2_ has left the site or formation of Q_B_^-^ before QH_2_ has cleared the channel and remains in the vicinity of the heme” (Cardona et al., 2012). Indeed, we observed different kinetics in electron transfer rates between initial and successive light exposures (**Figure. 2e**,**h**). Meaning that the “photosynthetic control” triggered changes in the output of electrons from PSII, which then pose limitations on the linear electron transfer rate and causes donor side limitations on PSI, resulting in a lower downstream yield.

Under anoxia, the plastoquinone/ol pool is over reduced since its major oxidizer, PTOX, is inactivated due to the lack of O_2_. Therefore, decreased activity of Cyt*b*_*6*_*f* would result in continuous limitations on PSII. However, under darkness, the activity of ATPase lowers the redox potential, which can also be seen by the appearance of a “phase b” rise in the kinetics of the charge separation (**Figure. 5c**). In addition, we do see that during short dark exposure (up to 5 minutes), the redox potential recovers its original low state, yet the posed conformational changes that enhance PSI cyclic electron flow and diminishes PSII linear electron flow remain (up to 15 minutes). Interestingly, the presence of DCMU did not result in any decrease in “phase a” of the measured ECS (**Supp. 5**), nor did it decrease the formation of a redox potential (**Figures. 5d**). However, the dissipation rate of the redox potential was increased, and following less than two minutes regained its initial low state. Taken together, these observations suggest that although the initiation of the “photosynthetic control” might not be related to the reduction state of the plastoquinone\ol pool, its dissipation is.

Another interesting theory, hypothesized by Prášil and coworkers (Prášil et al., 1996) postulated that when the photosynthetic electron transport chain is over-reduced, for an instance upon a dark-to-light, there is a decoupling between PSII photochemical yield and O_2_ evolution. Namely, PSII fluorescence yield recovered remarkably faster from pre-illumination than O_2_ evolution, in a manner that was dependent upon the reduction level of the plastoquinone pool. They also suggested that it was not due to a direct radiative charge back-reaction, but rather to the emergence of cyclic electron flow in PSII presumably via cytochrome *b559* (Prášil et al., 1996). This could explain the observed decrease in OEC activity which was observed in the TL measurements (**Figure. 5f**), and is also in line with the shifts in Chl*a* fluorescence kinetics (**Figure. 3c**), which show alternated rise of its trace at the onset of light. In any case, these processes would result in a decreased linear electron flow and assist the cells in subsiding oxidative damage, which could otherwise harm the viability of the cells. Yet, unfortunately, this decrease in the apparent linear electron flow also diminish the productivity of the photosynthetic apparatus, posing new barriers to cope with in the search for an improved energy yield via photosynthesis.

## Material & Methods

### Cells; type, growth, and conditions

The *Chlamydomonas reinhardtii* strain CC124 (137c mt-) was obtained from the *Chlamydomonas* Resource Center. *Chlorella sorokiniana* was a kind gift from the Nathan Nelson lab in Tel-Aviv University, Israel. In addition, we isolated a new *Monorophidium* strain from the Yarkon River in Israel and was identified it by PCR (see **Supp. 7**). All cultures were maintained and cultivated in Erlenmeyer flasks under continuous illumination (at an irradiance of 60 μE m^-2^ sec^-1^) in TAP or TP medium. Cell samples were taken from cultures in early log phase (between 2-5 mg Chl mL^-1^). Chlorophyll concentration was determined using 90% acetone or 100% methanol according to (Ritchie, 2006).

### Cell preparation for photosynthetic measurements

The cells were centrifuged to a final concentration of 20 mg Chl mL^-1^ in TAP (or TP for autotrophic cultures) +50mM HEPES, pH 7.2, and placed in a sealed quartz cuvette (5 mL). When indicated, Glucose oxidase (200 units mL^-1^), catalase (200 units mL^-1^), and Glucose (50 mM) were added to scavenge traces of O_2_. Anaerobic induction was achieved by keeping the cells in darkness for an hour. If indicated, 40 µM (final concentration) of 3-(3,4-dichlorophenyl)-1,1-dimethylurea (DCMU, Sigma-Aldrich) and 1mM (final concentration) of hydroxylamine (HA, Sigma-Aldrich) were added 10 minutes prior to the measurements. In all the experiments that were conducted in the JTS, 10% of Ficoll was added prior to induction, to avoid cell sedimentation.

### Combined gas exchange and photosynthetic efficiency measurements

For tracking the concentration of diffused gasses, such as; H_2_, O_2_ and CO_2_, the experimental cuvette was placed in a home-built membrane inlet mass spectrometer (MIMS), as described in (Milrad et al., 2021). Simultaneously, NADPH and Chlorophyll *a* fluorescence measurement were conducted using a DUAL-PAM-100 (Heinz Walz Gmbh, Effeltrich, Germany). Supplied with NADPH/9-AA emitter/detector module as described in (Milrad et al., 2021; Schreiber and Klughammer, 2009). Monitoring state transition was conducted by taking cell samples directly from the experimental cuvette (50 μL) and injecting them into a glass capillary, and inserting it to a glass Dewar (Horiba) filled with liquid nitrogen (at 77°K). Samples were then measured in a fluorimeter (Horiba Fluoromax-4), with excitation light was of 435 nm. Emission was measured between 650-750 nm.

### Spectroscopic analysis of the redox potential and electron flux

Electrochromic shifts (ECS) were detected using a Joliot Type Spectrometer (JTS-100, Biologic SAS, France), supplied with a BiLED, which is able to simultaneously measure the absorbance of 520 and 546 nm, as described in (Milrad et al., 2021). When indicated, laser flashes were pumped by a frequency-doubled Nd:YAG laser (Litron nano), excitation wavelength was adjusted using a dye (DCM, exciton laser dye). When cells are exposed to a single turnover laser flash their absorbance is changed via a 3-phase shift (Bailleul et al., 2010). The initial increase in the ECS signal (termed “phase a”) lasts less than a millisecond and is a product of charge separations taking place in both photosystems, hence decreased signal can report on their degradation. During the second phase (termed “phase b”),which usually lasts up to 10 milliseconds (Buchert et al., 2020), ECS signal increases due to the build-up of membrane potential, mainly via Cyt*b*_*6*_*f*. In the final phase (termed “phase c”), the signal decreases exponentially due to the breakup dissipation of the membrane potential via ATPase activity. Since no differences in the amplitude of “phase a” were observed (**Supp. 3**), the results were normalized it. Observed differences in the kinetics of both the apparent “phase b” and “phase c” were used to determine shifts in the redox potential across the thylakoid membrane. Dark interval relaxation kinetics (DIRK) was measured during illumination (orange ring, 630 nm) by exposing the cells to short dark pulses (60 msec), and subtracting the steady state kinetics (which did not show much drift) from the rate of the decrease in the signal during darkness. The rate was then normalized to “phase a” which was triggered by the laser flash prior to illumination. Since we have observed no decrease in ECS due to the addition of DCMU, the results are not correlated to PSI but rather to general PS, although they resulted in the same apparent values.

### Photosystem I reduction state analysis

PSI efficiency was also measured in the JTS, using a BiLED, which can simultaneously measure the absorbance of 705 and 740 nm. Following dark anaerobic induction, the cells were exposed to two light fluctuations (at an irradiance of 220 µE m ^−2^ sec ^−1^, Ps’), with three minutes of darkness in between them. During each illumination, the cells were exposed to a short high light pulse (30 msec at an irradiance of 2000 µE m ^−2^ sec ^−1^, Pm’), followed by a dark pulse (for 4 sec, **Figure. 3**; gray background, Po’). These “high light/Dark” pulses were repeated every 15 seconds for a total duration of two minutes, and each pulse was shifted in accordance with the value of Po’ at the end of its dark period. Results were analyzed according to (Buchert et al., 2020).

### Thermoluminescence measurements

The cells were centrifuged shortly, transferred to a minimal medium (TP) in a final concentration of 15 mg Chl mL^-1^. Aerobic samples were dark-adapted for 20 minutes before measurements, while anaerobic cultures were placed in a 13 mL hypo-vial bottles, and flushed with N_2_ for 10 minutes under darkness, followed by an additional hour of dark-incubation with continuous shaking. Illuminated cultures were subjected to two minutes of light exposure (at an irradiance of 370 μE m^-2^ sec^-1^), followed by either 5 or 15 minutes of darkness before they were sampled (300 µL). TL measurements were carried out by a custom-made TL apparatus, as described in (Podmaniczki et al., 2020). For the illumination with STF and the recording of the TL signal, the samples were placed on a copper plate in air, connected to a cold finger immersed in liquid N_2_. A heater coil (SEI 10/50, Thermocoax, France) ensured the desired temperature of the sample during the measurement. It should be noted that in the case of whole cells, freezing before TL measurements is not recommended due to possible cellular damage (Ducruet, 2013), thus algal samples were illuminated by two single turnover saturating Xe flashes (of 1.5 μsec duration at half-peak intensity), at 4 °C, and the sample was heated to 70°C in darkness with a heating rate of 20°C min^-1^. The emitted TL was measured with a photomultiplier (H10721-20, Hamamatsu, Japan) simultaneously with recording the temperature.

## Acknowledgements

We thank M. Hippler and F. Buchert from WWU Münster, Germany for constructive discussion. We thank Tamar Elman and N. Shahar from TAU, Israel for carefully reading the manuscript and for their comments.

## References

Allen, J.F. (2003). Cyclic, pseudocyclic and noncyclic photophosphorylation: New links in the chain. Trends Plant Sci. 8, 15–19.

Alric, J., Lavergne, J., and Rappaport, F. (2010). Redox and ATP control of photosynthetic cyclic electron fl ow in Chlamydomonas reinhardtii (I) aerobic conditions. Biochim. Biophys. Acta 1797, 44–51.

Antal, T.K., Petrova, E., Slepnyova, V., Kukarskikh, G., Volgusheva, A.A., Dubini, A., Baizhumanov, A., Tyystjärvi, T., Gorelova, O., Baulina, O., et al. (2020). Photosynthetic hydrogen production as acclimation mechanism in nutrient-deprived Chlamydomonas. Algal Res. 49, 101951.

Bailleul, B., Cardol, P., Breyton, C., and Finazzi, G. (2010). Electrochromism: A useful probe to study algal photosynthesis. Photosynth. Res. 106, 179–189.

Brinkert, K., De Causmaecker, S., Krieger-Liszkay, A., Fantuzzi, A., and Rutherford, A.W. (2016). Bicarbonate-induced redox tuning in Photosystem II for regulation and protection. Proc. Natl. Acad. Sci. U. S. A. 113, 12144–12149.

Buchert, F., Mosebach, L., Gäbelein, P., and Hippler, M. (2020). PGR5 is required for efficient Q cycle in the cytochrome b6f complex during cyclic electron flow. Biochem. J. 477, 1631–1650.

Burlacot, A., Sawyer, A.L., Cuiné, S., Auroy-Tarrago, P., Blangy, S., Happe, T., and Peltier, G. (2018). Flavodiiron-mediated O2 photoreduction links H2 production with CO2 fixation during the anaerobic induction of photosynthesis. Plant Physiol. 177, 1639–1649.

Burlacot, A., Peltier, G., and Li-beisson, Y. (2019). Subcellular Energetics and Carbon Storage in Chlamydomonas. Cells 8, 1–15.

Cardona, T., Sedoud, A., Cox, N., and Rutherford, A.W. (2012). Charge separation in Photosystem II: A comparative and evolutionary overview. Biochim. Biophys. Acta - Bioenerg. 1817, 26–43.

Chaux, F., Burlacot, A., Mekhalfi, M., Auroy, P., Blangy, S., Richaud, P., and Peltier, G. (2017). Flavodiiron proteins promote fast and transient O2 photoreduction in chlamydomonas. Plant Physiol. 174, 1825–1836.

Ducruet, J.M. (2013). Pitfalls, artefacts and open questions in chlorophyll thermoluminescence of leaves or algal cells. Photosynth. Res. 115, 89–99.

Ducruet, J.M., and Vass, I. (2009). Thermoluminescence: Experimental. Photosynth. Res. 101, 195–204.

Dufková, K., Barták, M., Morkusová, J., Elster, J., and Hájek, J. (2019). Screening of growth phases of antarctic algae and cyanobacteria cultivated on agar plates by chlorophyll fluorescence imaging. Czech Polar Reports 9, 170–181.

Ghysels, B., Godaux, D., Matagne, F.R., Cardol, P., and Franck, F. (2013). Function of the Chloroplast Hydrogenase in the Microalga Chlamydomonas: The Role of Hydrogenase and State Transitions during Photosynthetic Activation in Anaerobiosis. Public Libr. Sci. 8.

Godaux, D., Bailleul, B., Berne, N., and Cardol, P. (2015). Induction of Photosynthetic Carbon Fixation in Anoxia Relies on Hydrogenase Activity and Proton-Gradient Regulation-Like1-Mediated Cyclic Electron Flow in Chlamydomonas reinhardtii 1. Plant Physiol. 168, 648–658.

Hemschemeier, A., and Happe, T. (2011). Alternative photosynthetic electron transport pathways during anaerobiosis in the green alga Chlamydomonas reinhardtii. Biochim. Biophys. Acta 1807, 919–926.

Houille-vernes, L., Rappaport, F., Wollman, F.-A., Alric, J., and Johnson, X. (2011). Plastid terminal oxidase 2 (PTOX2) is the major oxidase involved in chlororespiration in Chlamydomonas. Pnas 108, 20820–20825.

Jokel, M., Johnson, X., Peltier, G., Aro, E.M., and Allahverdiyeva, Y. (2018). Hunting the main player enabling Chlamydomonas reinhardtii growth under fluctuating light. Plant J. 822–835.

Jokel, M., Nagy, V., Tóth, S.Z., Kosourov, S.N., and Allahverdiyeva, Y. (2019). Elimination of the flavodiiron electron sink facilitates long-term H2 photoproduction in green algae. Biotechnol. Biofuels 12, 1–16.

Kaplan, A., and Reinhold, L. (1999). CO2 Concentrating Mechanisms in Microorganisms. Transport 50, 539–570.

Kirchhoff, H., Li, M., and Puthiyaveetil, S. (2017). Sublocalization of Cytochrome b6f Complexes in Photosynthetic Membranes. Trends Plant Sci. 22, 574–582.

Klughammer, C., and Schreiber, U. (1994). An improved method, using saturating light pulses, for the determination of photosystem I quantum yield via P700+-absorbance changes at 830 nm. Planta 192, 261–268.

Kodru, S., Malavath, T., Devadasu, E., Nellaepalli, S., Stirbet, A.D., Subramanyam, R., and Govindjee (2015). The slow S to M rise of chlorophyll a fluorescence reflects transition from state 2 to state 1 in the green alga Chlamydomonas reinhardtii. Photosynth. Res. 125, 219–231.

Koroidov, S., Shevela, D., Shutova, T., Samuelsson, G., and Messinger, J. (2014). Mobile hydrogen carbonate acts as proton acceptor in photosynthetic water oxidation. Proc. Natl. Acad. Sci. U. S. A. 111, 6299–6304.

Kosourov, S.N., Jokel, M., Aro, E.M., and Allahverdiyeva, Y. (2018). A new approach for sustained and efficient H2 photoproduction by Chlamydomonas reinhardtii †. Energy Environ. Sci. 11, 1431–1436.

Kosourov, S.N., Nagy, V., Shevela, D., Jokel, M., Messinger, J., and Allahverdiyeva, Y. (2020). Water oxidation by photosystem II is the primary source of electrons for sustained H2photoproduction in nutrient-replete green algae. Proc. Natl. Acad. Sci. U. S. A. 117, 29629–29636.

Kozuleva, M., Petrova, A., Milrad, Y., Semenov, A., Ivanov, B., Redding, K.E., and Yacoby, I. (2021). Phylloquinone is the principal Mehler reaction site within photosystem I in high light. Plant Physiol. 186, 1848–1858.

Krause, G.H. (1988). Photoinhibition of photosynthesis. An evaluation of damaging and protective mechanisms. Physiol. Plant. 74, 566–574.

Liran, O., Semyatich, R., Milrad, Y., Eilenberg, H., Weiner, I., and Yacoby, I. (2016). Microoxic Niches within the Thylakoid Stroma of Air-Grown Chlamydomonas reinhardtii Protect [FeFe] - Hydrogenase and Support Hydrogen Production under Fully Aerobic Environment. Plant Physiol. 172, 264–271.

Milrad, Y., Schweitzer, S., Feldman, Y., and Yacoby, I. (2018). Green algal hydrogenase activity is outcompeted by carbon fixation before inactivation by oxygen takes place. Plant Physiol. 177, 918–926.

Milrad, Y., Schweitzer, S., Feldman, Y., and Yacoby, I. (2021). Bi-directional electron transfer between H2 and NADPH mitigates light fluctuation responses in green algae. Plant Physiol. 186, 168–179.

Nawrocki, W.J., Tourasse, N.J., Taly, A., Rappaport, F., and Wollman, F.-A. (2015). The Plastid Terminal Oxidase: Its Elusive Function Points to Multiple Contributions to Plastid Physiology. Annu. Rev. Plant Biol. 66, 49–74.

Peloquin, J.M., and Britt, R.D. (2001). EPR/ENDOR characterization of the physical and electronic structure of the OEC Mn cluster. Biochim. Biophys. Acta - Bioenerg. 1503, 96–111.

Podmaniczki, A., Nagy, V., Vidal-Meireles, A., Tóth, D., Patai, R., Kovács, L., and Tóth, S.Z. (2020). Ascorbate inactivates the oxygen-evolving complex in prolonged darkness. Physiol. Plant. 1–14.

Prášil, O., Kolber, Z., Berry, J.A., and Falkowski, P.G. (1996). Cyclic electron flow around Photosystem II in vivo. Photosynth. Res. 395–410.

Ritchie, R.J. (2006). Consistent sets of spectrophotometric chlorophyll equations for acetone, methanol and ethanol solvents. Photosynth. Res. 89, 27–41.

Roberty, S., Bailleul, B., Berne, N., Franck, F., and Cardol, P. (2014). PSI Mehler reaction is the main alternative photosynthetic electron pathway in Symbiodinium sp., symbiotic dinoflagellates of cnidarians. New Phytol. 204, 81–91.

Sacksteder, C.A., and Kramer, D.M. (2000). Dark-interval relaxation kinetics (DIRK) of absorbance changes as a quantitative probe of steady-state electron transfer. Photosynth. Res. 66, 145–158.

Saito, K., Rutherford, A.W., and Ishikita, H. (2013). Mechanism of proton-coupled quinone reduction in Photosystem II. Proc. Natl. Acad. Sci. U. S. A. 110, 954–959.

Schöttler, M.A., and Tóth, S.Z. (2014). Photosynthetic complex stoichiometry dynamics in higher plants: Environmental acclimation and photosynthetic flux control. Front. Plant Sci. 5, 1–15.

Schöttler, M.A., Tóth, S.Z., Boulouis, A., and Kahlau, S. (2015). Photosynthetic complex stoichiometry dynamics in higher plants: Biogenesis, function, and turnover of ATP synthase and the cytochrome b6f complex. J. Exp. Bot. 66, 2373–2400.

Schreiber, U., and Klughammer, C. (2009). New NADPH/9-AA module for the DUAL-PAM-100: Description, operation and examples of application. PAM Appl. Notes 1–13.

Schreiber, U., Hormann, H., Neubauer, C., and Klughammer, C. (1995). Assessment of Photosystem II Photochemical Quantum Yield by Chlorophyll Fluorescence Quenching Analysis. Plant Physiol. 22, 209–220.

Shevela, D., Do, H.N., Fantuzzi, A., Rutherford, A.W., and Messinger, J. (2020). Bicarbonate-Mediated CO2 Formation on Both Sides of Photosystem II. Biochemistry 59, 2442–2449.

Sipka, G., Magyar, M., Mezzetti, A., Akhtar, P., Zhu, Q., Xiao, Y., Han, G., Santabarbara, S., Shen, J.-R., Lambrev, P.H., et al. (2021). Light-adapted charge-separated state of photosystem II: structural and functional dynamics of the closed reaction center. Plant Cell 33, 1286–1302.

Stirbet, A.D., and Govindjee (2011). On the relation between the Kautsky effect (chlorophyll a fluorescence induction) and Photosystem II: Basics and applications of the OJIP fluorescence transient. J. Photochem. Photobiol.

Strasserf, R.J., Srivastava, A., and Govindjee (1995). Polyphasic chlorophyll a fluorescence transient in plants and cyanobacteria. Photochem. Photobiol. 61, 32–42.

Stripp, S.T., Goldet, G., Brandmayr, C., Sanganas, O., Vincent, K.A., Haumann, M., Armstrong, F.A., and Happe, T. (2009). How oxygen attacks [FeFe] hydrogenases from photosynthetic organisms. Proc. Natl. Acad. Sci. 106, 1–6.

Takahashi, H., Clowez, S., Wollman, F.-A., Vallon, O., and Rappaport, F. (2013). Cyclic electron flow is redox-controlled but independent of state transition. Nat. Comun.

Tikhonov, K., Shevela, D., Klimov, V. V., and Messinger, J. (2018). Quantification of bound bicarbonate in photosystem II. Photosynthetica 56, 210–216.

Tolleter, D., Ghysels, B., Alric, J., Petroutsos, D., Tolstygina, I., Krawietz, D., Happe, T., Auroy, P., Adriano, J.-M., Beyly-Adriano, A., et al. (2011). Control of Hydrogen Photoproduction by the Proton Gradient Generated by Cyclic Electron Flow in Chlamydomonas reinhardtii. Plant Cell 23, 2619–2630.

Umena, Y., Kawakami, K., Shen, J.R., and Kamiya, N. (2011). Crystal structure of oxygen-evolving photosystem II at a resolution of 1.9Å. Nature 473, 55–60.

Vass, I., and Cser, K. (2009). Janus-faced charge recombinations in photosystem II photoinhibition. Trends Plant Sci. 14, 200–205.

Vinyard, D.J., Ananyev, G.M., and Dismukes, G.C. (2013). Photosystem II: The Reaction Center of Oxygenic. Annu. Rev. Biochem. 82, 577–606.

