## Supplementary material for "A novel PSII photosynthetic control is activated in anoxic cultures of green algae": Supp

### Supplemental data

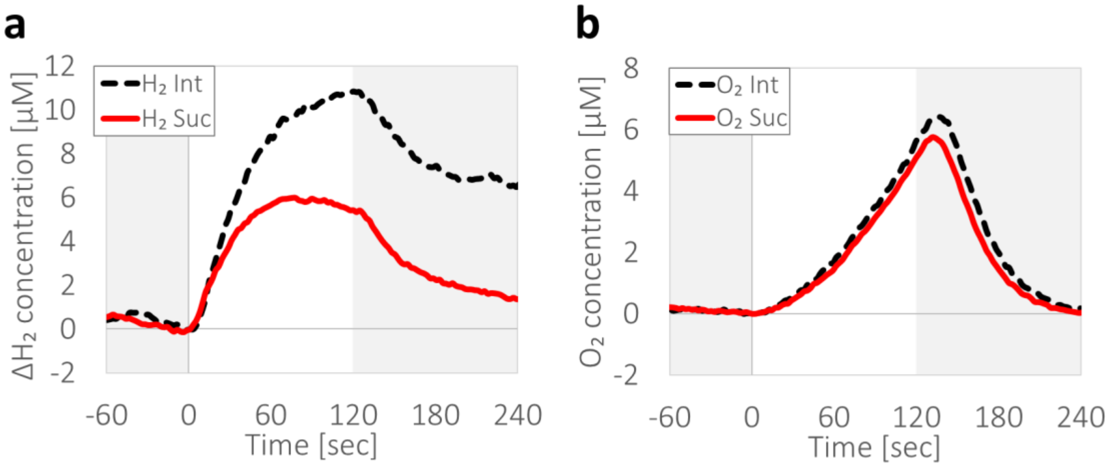


**Supplemental Figure 1: Hydrogen and oxygen evolution under autotrophic conditions.** *C. reinhardtii* (wild-type strain- CC124) cells were cultivated under autotrophic conditions (TP, pH 7.2, supplied with 5% $CO_2$) and incubated for an hour under dark anaerobiosis, after which they were challenged with light fluctuations of 2 minutes under illumination (at an irradiance of 370 µE $m^{-2}$ $s^{-1}$, white background), followed by 3 minutes of darkness (gray background, as shown in Figure 1). Shown are the differences between the initial light exposure (dashed black) and the average of the successive exposures (solid red). $H_2$ (**a**) and $O_2$ (**b**) concentrations were measured. Graphs represent an average of three biological repeats.

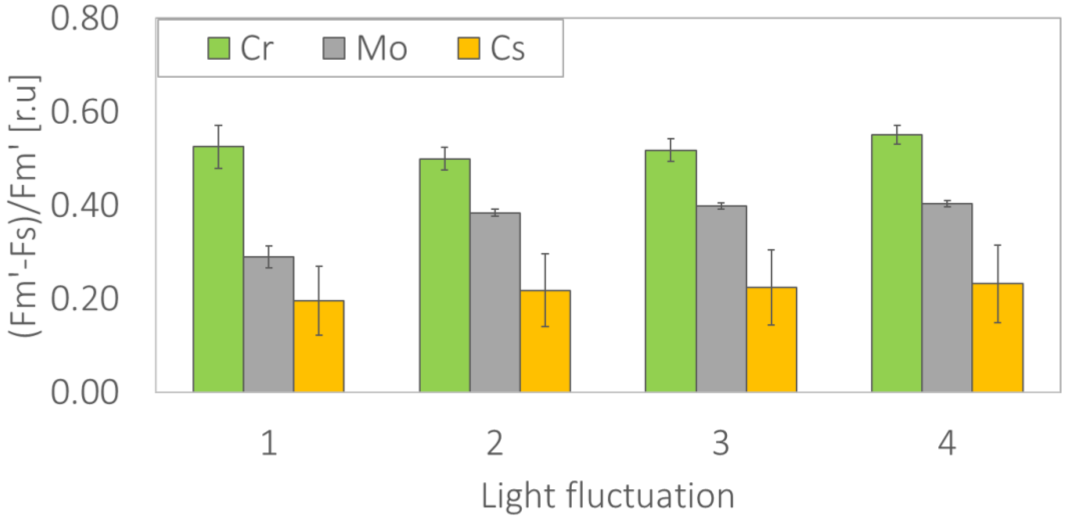


**Supplemental Figure 2: Quantum yield under light fluctuations.** Following an hour of dark anaerobic incubation, three algae species: *Chlamydomonas reinhardtii* (Cr, green), a newly isolated strain of *Monoraphidium* (Mo, gray), and *Chlorella sorokiniana* (Cs, yellow) were examined in a Dual-PAM-100 for Chl*a* fluorescence under fluctuating light exposures. During illuminations, the cells were also exposed to saturating light pulses (see yellow arrows in **Figure**

**1a**) to evaluate the maximal fluorescence (Fm'). Relative quantum yields were calculated as (Fm'-Fs)/Fm' for each fluctuation and compared. Each column represent the averaged result of at least three biological repeats. Error bars represent standard error.

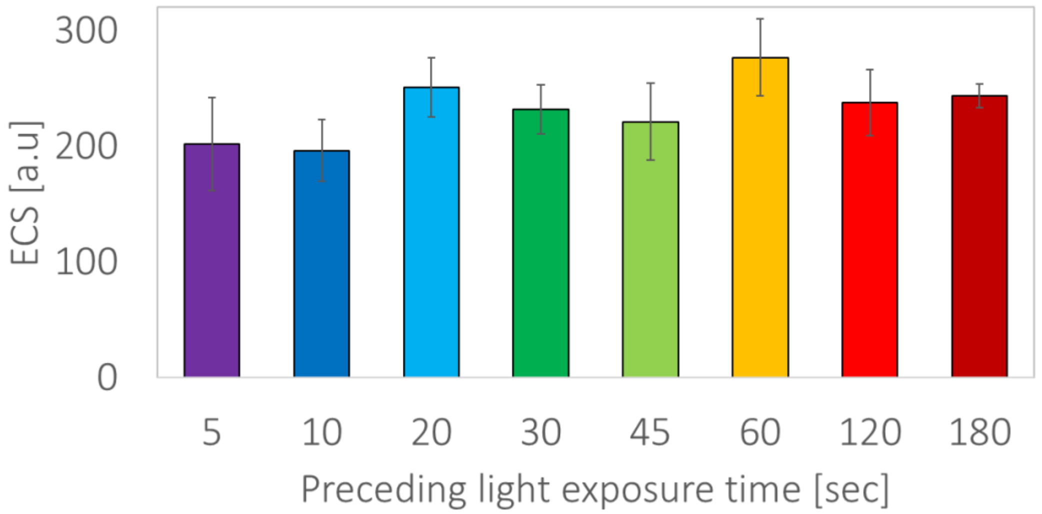


**Supplemental Figure 3: Charge separation of laser flashes following illumination.** *C. reinhardtii* wild-type strain CC124 cells were incubated for an hour under dark anaerobiosis, after which they tested in JTS-100, equipped with a BiLED (520-546 nm) measuring lamp. The cells were exposed to illumination (at an irradiance of 370 μE $m^{-2}$ $s^{-1}$) for a duration of either 5, 10, 20, 30, 45, 60, 120 or 180 seconds, after which they were exposed to a 5 ns laser flash. Presented are the values for "phase a" of the charge separation, *i.e* the differences in ECS following less than a millisecond. Columns represent the average of at least three biological repeats. Error bars represent standard error.

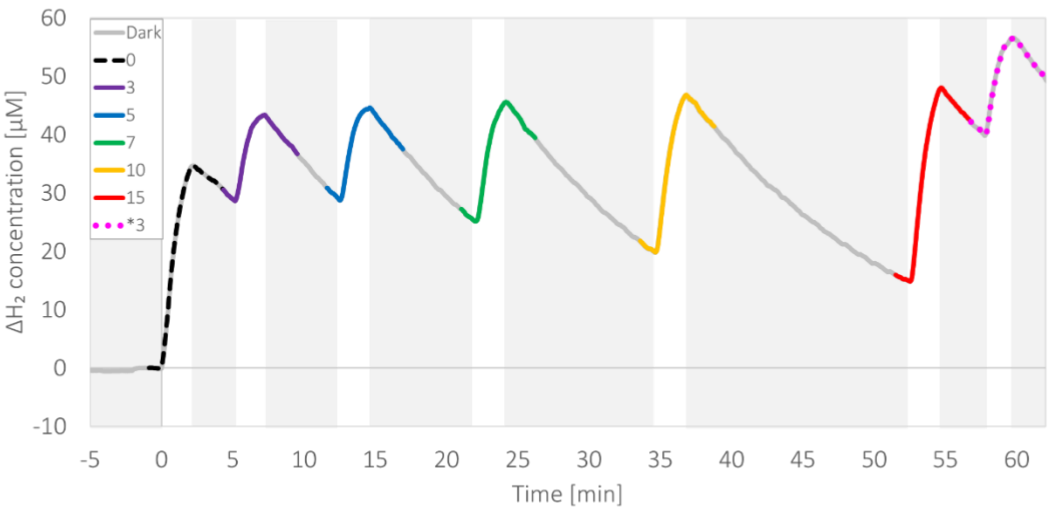

**Supplemental Figure 4: Darkness reestablish the initial fast electron flux.** Following an hour of dark incubation in the presence of $O_2$ scavengers (GOx), *C. reinhardtii* wild-type strain CC124 cells were subjected to a series of fixed duration of light exposures (370 µE $m^{-2}$ $s^{-1}$ for 2 minutes, white background) hatched with an increasing duration of dark incubations 0-15 minutes. (**a**) $H_2$ accumulation as a function of the preceded dark incubation time (gray background) between fixed 2 minutes of exposures to light. The evolution of $H_2$ in each light exposure was measured by MIMS and the trace is highlighted according to its preceding duration of dark incubation (initial exposure, 0 – dashed, 3 min – purple, 5 min – blue, 7 min – green, 10 min – yellow, 15 min – red, additional 3 min – dotted pink, same color index was used for all the traces in all panels). Each experiment was repeated using at least three biological replicates. Error bars indicate standard error ($n \geq 3$).

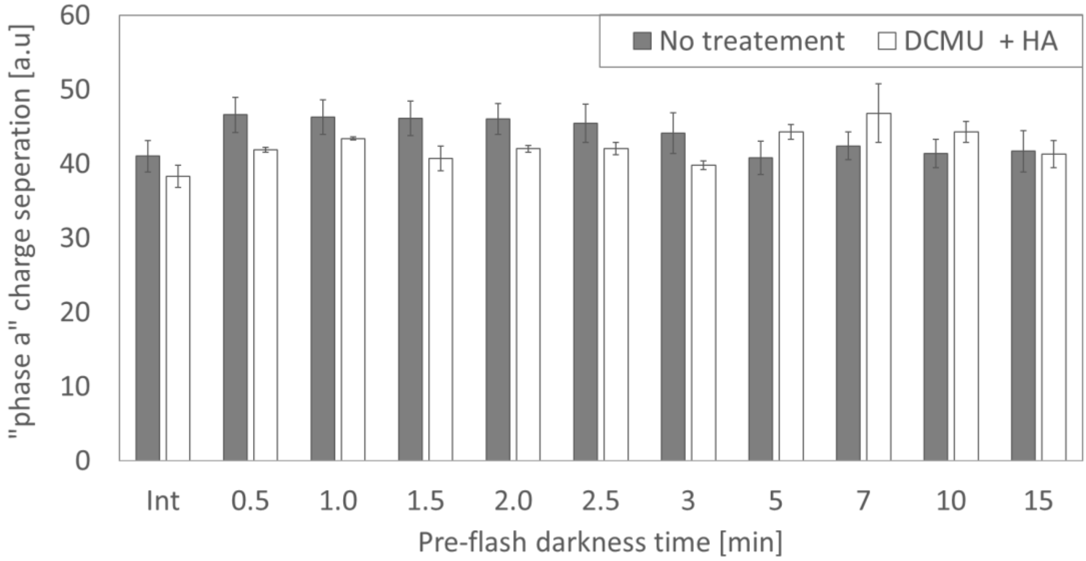


**Supplemental Figure 5: Charge separation of laser flashes following prolonged darkness.** *C. reinhardtii* wild-type strain CC124 cells were incubated for an hour under dark anaerobiosis in the absence (gray) or presence (white) of DCMU and hydroxylamine (HA), after which they tested in JTS-100, equipped with a BiLED (520-546 nm) measuring lamp. The cells were then exposed to a 5 ns laser flash (int- dashed black), illuminated for two minutes (at an irradiance of 370 µE $m^{-2}$ $s^{-1}$), and then exposed again to laser flashes (as specified for each column). Presented are the values for "phase a" of the charge separation, *i.e* the differences in ECS following less than a millisecond. Columns represent the average of at least three biological repeats. Error bars represent standard error.

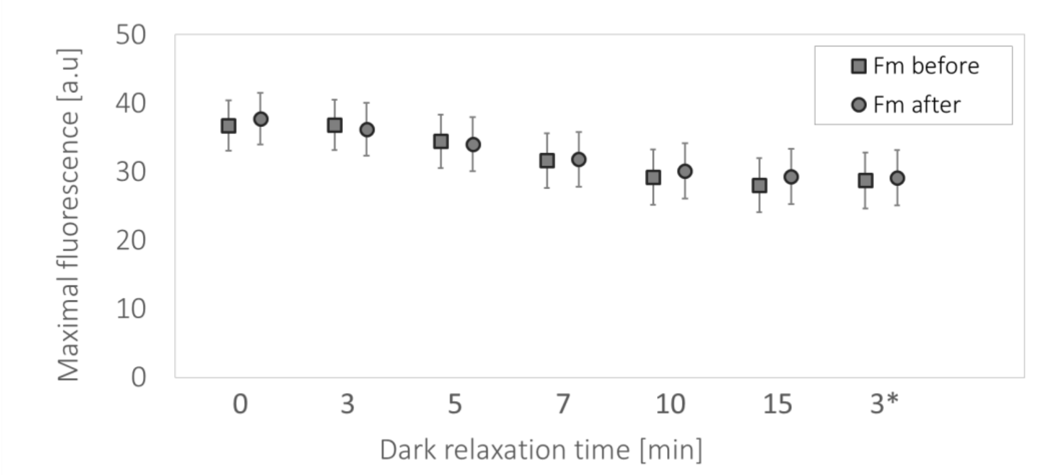


**Supplemented Figure. 6: Maximal Chl*a* fluorescence differences as a function of the duration of light exposure.** Following an hour of dark incubation in the presence of $O_2$ scavengers (GOx), *C. reinhardtii* wild-type strain CC124 cells were subjected to a series of light exposures (370 µE $m^{-2}$ $s^{-1}$ for 2 minutes), in which dark relaxation time between exposures was gradually increased (X-Axis). 60 seconds before (square) and 30 seconds after (circle) each illumination, the cells were subjected to a saturating pulse to determine maximal Chl*a* fluorescence (Fm). Error bars indicate standard error ($n \geq 3$).

| Select for downloading or viewing reports | Description | Scientific Name | Max Score | Total Score | Query Cover | E value | Per. Ident | Acc. Len | Accession |
|---|---|---|---|---|---|---|---|---|---|
| Select seq KT180321.1 | Monoraphidium sp. SDEC-17 18S ribosomal RNA gene, partial sequence | Monoraphidium sp. SDEC-17 | 1773 | 1773 | 88% | 0.0 | 96.33% | 1705 | KT180321.1 |
| Select seq MW683514.1 | Monoraphidium sp. EHY small subunit ribosomal RNA gene, partial sequence | Monoraphidium sp. EHY | 1768 | 1768 | 88% | 0.0 | 96.24% | 1713 | MW683514.1 |
| Select seq MG022711.1 | Monoraphidium convolutum isolate CCAP 202/10F small subunit ribosomal RNA gene, partial sequence | Monoraphidium convolutum | 1766 | 1766 | 89% | 0.0 | 95.83% | 1267 | MG022711.1 |
| Select seq MH340049.1 | Monoraphidium sp. HDMA-11 18S small subunit ribosomal RNA gene, partial sequence | Monoraphidium sp. HDMA-11 | 1762 | 1762 | 88% | 0.0 | 96.14% | 1773 | MH340049.1 |
| Select seq KT833590.1 | Monoraphidium contortum voucher CCMA UFSCar 473 18S ribosomal RNA gene, partial sequence | Monoraphidium contortum | 1762 | 1762 | 88% | 0.0 | 96.14% | 2128 | KT833590.1 |
| Select seq KM199735.1 | Monoraphidium sp. QLY-1 18S ribosomal RNA gene, partial sequence | Monoraphidium sp. QLY-1 | 1762 | 1762 | 88% | 0.0 | 96.14% | 1680 | KM199735.1 |
| Select seq MW507790.1 | Monoraphidium sp. SP03 small subunit ribosomal RNA gene, partial sequence | Monoraphidium sp. SP03 | 1762 | 1762 | 88% | 0.0 | 96.14% | 1218 | MW507790.1 |
| Select seq JQ809706.1 | Monoraphidium sp. FXY-10 18S ribosomal RNA gene, partial sequence | Monoraphidium sp. FXY-10 | 1762 | 1762 | 88% | 0.0 | 96.14% | 1730 | JQ809706.1 |
| Select seq HM483518.1 | Rhombocystis complanata strain KR 1998/2 18S ribosomal RNA gene, partial sequence | Rhombocystis complanata | 1762 | 1762 | 88% | 0.0 | 96.14% | 2877 | HM483518.1 |
| Select seq KX671912.1 | Monoraphidium sp. B LP-2016 18S ribosomal RNA gene, partial sequence | Monoraphidium sp. B LP-2016 | 1760 | 1760 | 89% | 0.0 | 95.74% | 2466 | KX671912.1 |
| Select seq KJ616758.1 | Monoraphidium dybowskii strain LB53 18S ribosomal RNA gene, partial sequence | Monoraphidium dybowskii | 1759 | 1759 | 88% | 0.0 | 96.05% | 1787 | KJ616758.1 |
| Select seq KT833596.1 | Messastrum gracile voucher CB 2009/3 18S ribosomal RNA gene, partial sequence | | | | | | | | |

**Supplemented Figure. 7: PCR results for culture isolation**. Water samples were taken from the Yarkon River in Israel. Algal cultures were grown under autotrophic conditions under illumination in the presence of antimycin to reduce bacterial contamination. Single colonies were then taken for PCR determination. The results indicate that the cells are indeed from the species *Monoraphidium*. 7
